# Phosphatidylserine prevents the generation of a protein-free giant plasma membrane domain in yeast

**DOI:** 10.1101/2020.08.10.245530

**Authors:** Tetsuo Mioka, Guo Tian, Wang Shiyao, Takuma Tsuji, Takuma Kishimoto, Toyoshi Fujimoto, Kazuma Tanaka

**Author notes:** Corresponding author: Tetsuo Mioka and Kazuma Tanaka, (TM) and (KT).

## Abstract

Membrane phase separation accompanied with micron-scale domains of lipids and proteins occurs in artificial membranes; however, a similar large phase separation has not been reported in the plasma membrane of the living cells. We demonstrate here that a stable micron-scale protein-free region is generated in the plasma membrane of the yeast mutants lacking phosphatidylserine. We named this region the “void zone”. Transmembrane proteins, peripheral membrane proteins, and certain phospholipids are excluded from the void zone. The void zone is rich in ergosterol and requires ergosterol and sphingolipids for its formation. These characteristics of the void zone are similar to the properties of the cholesterol-enriched domain in phase-separated artificial membranes. We propose that phosphatidylserine prevents the formation of the void zone by preferentially interacting with ergosterol. We also found that void zones were frequently in contact with vacuoles, in which a membrane domain was also formed at the contact site.

**Summary statement:** Yeast cells lacking phosphatidylserine generate protein-free plasma membrane domains, and vacuoles contact with this domain. This is the first report of micron-scale plasma membrane domains in living cells.

## Introduction

The fluid mosaic model describing the dynamic distribution of proteins at the plasma membrane has been largely modified and developed to date (Singer and Nicolson, 1974; Nicolson, 2014; Kusumi A *et al*., 2012). Lateral diffusion of proteins is not free and is influenced by protein interaction with other plasma membrane proteins and cytoskeletal elements. In cholesterol-rich domains, such as lipid rafts, certain proteins can accumulate due to protein-protein or protein-lipid interactions (Lingwood and Simons, 2010). The plasma membrane is currently considered to be a nanoscale heterogeneous structure. In addition, several macroscopic diffusion barriers have been detected, and some of the barriers represent membrane compartmentalization due to interactions between the cytoskeleton and membrane proteins (Kusumi *et al*., 2012; Trimble and Grinstein, 2015). In artificial membranes, such as giant unilamellar vesicles (GUVs) and giant plasma membrane vesicles (GPMVs), membrane phase separation leads to the formation of even larger domains of proteins and lipids (Veatch and Keller, 2003; Baumgart *et al*., 2007; Elson *et al*., 2010; Carquin *et al*., 2016). Phase separation in artificial membranes has been well studied and is often compared to the nanoscale membrane domains found in the cells; however, large-scale phase separation is not observed in the plasma membranes of the living cells due to unknown reasons.

The plasma membranes are composed of diverse lipid species, and the role of phosphatidylserine (PS) and phosphatidylinositol phosphates (PIPs) in various cellular functions has been studied (Uchida *et al*., 2011; Cho *et al*., 2015; Middel *et al*., 2016; Tsuchiya *et al*., 2018; Michell, 2008; Balla, 2013). However, it is poorly understood how individual phospholipids influence the membrane environment.

Although PS is essential for growth of mammalian cells, yeast mutant cells lacking *CHO1*, the only PS synthase in the budding yeast, can grow (Arikketh *et al*., 2008; Atkinson *et al*., 1980). To explore a new role for PS, we have analysed PS-deficient *cho1*Δ yeast cells. In this study, we show that stable large protein-free membrane domains are detected in the plasma membrane of PS-deficient *cho1*Δ cells, which we named the “void zone”. Transmembrane proteins, peripheral membrane proteins, and certain phospholipids are excluded from the void zone. This property is very similar to the cholesterol-enriched membrane domain in phase-separated artificial membranes. Our results suggest that PS suppresses the development of large-scale phase separation in the plasma membrane of the living cells and consequently ensures the distribution of proteins and lipids throughout the plasma membrane. Furthermore, we found that vacuoles, the lysosomal organelle of yeast, contact with the void zone on the plasma membrane.

## Results

### PS-deficient cells show a protein-free region, “void zone”, in the plasma membrane

GFP-Snc1-pm, a mutant of v-SNARE Snc1, is uniformly distributed throughout the plasma membrane due to a defect in its endocytosis (Lewis *et al*. 2000). In PS-deficient *cho1*Δ cells grown at 37°C, GFP-Snc1-pm was heterogeneously distributed on the plasma membrane (Figure 1A). A GFP-Snc1-pm-deficient region was barely detectable at 30°C, was frequently present during incubation at 37°C for over 6 hours, and was not detected after heat shock at 42°C for 20 min (Figure 1B). This Snc1-pm-deficient region of the plasma membrane is referred to as “void zone” in the present study. The shape of the void zone observed on the cell surface was irregular and did not correspond to a smooth circle, and some cells had multiple void zones (Figure 1C). When *cho1*Δ cells were observed immediately after staining with FM4-64 lipophilic dye, FM4-64 was distributed throughout the plasma membrane including the void zone (Figure 1D), suggesting that the plasma membrane is not lost or significantly damaged in cells harbouring the void zone. To examine whether the void zone influences the distribution of other transmembrane proteins, four different transmembrane proteins, Pma1, Pdr5, Pdr12, and Sfk1, were compared with Snc1-pm (Figure 1E). Pma1 is the major plasma membrane H^+^-ATPase (Serrano *et al*., 1986). Pdr5 and Pdr12 are the ATP-binding cassette (ABC) transporters involved in the multidrug resistance and the weak organic acid resistance, respectively (Bauer *et al*., 1999). Sfk1 regulates phospholipid asymmetry in conjunction with the flippase complex Lem3-Dnf1/2 and is involved in proper localization of a phosphatidylinositol-4-kinase Stt4 (Audhya and Emr, 2002; Mioka *et al*., 2018). The results indicate that all these proteins showed void zones in the same region with void zones detected by Snc1-pm; the percentage of overlapping void zones was more than 82% in all cases (Figure 1E; percentage was determined from 70-110 cells with void zones). These results suggest that the void zone is a membrane protein-free region common to all other membrane proteins (see below), and the void zone can represent an abnormal lipid domain that inhibits the lateral movement of the transmembrane proteins into the domain.

**Figure 1.**
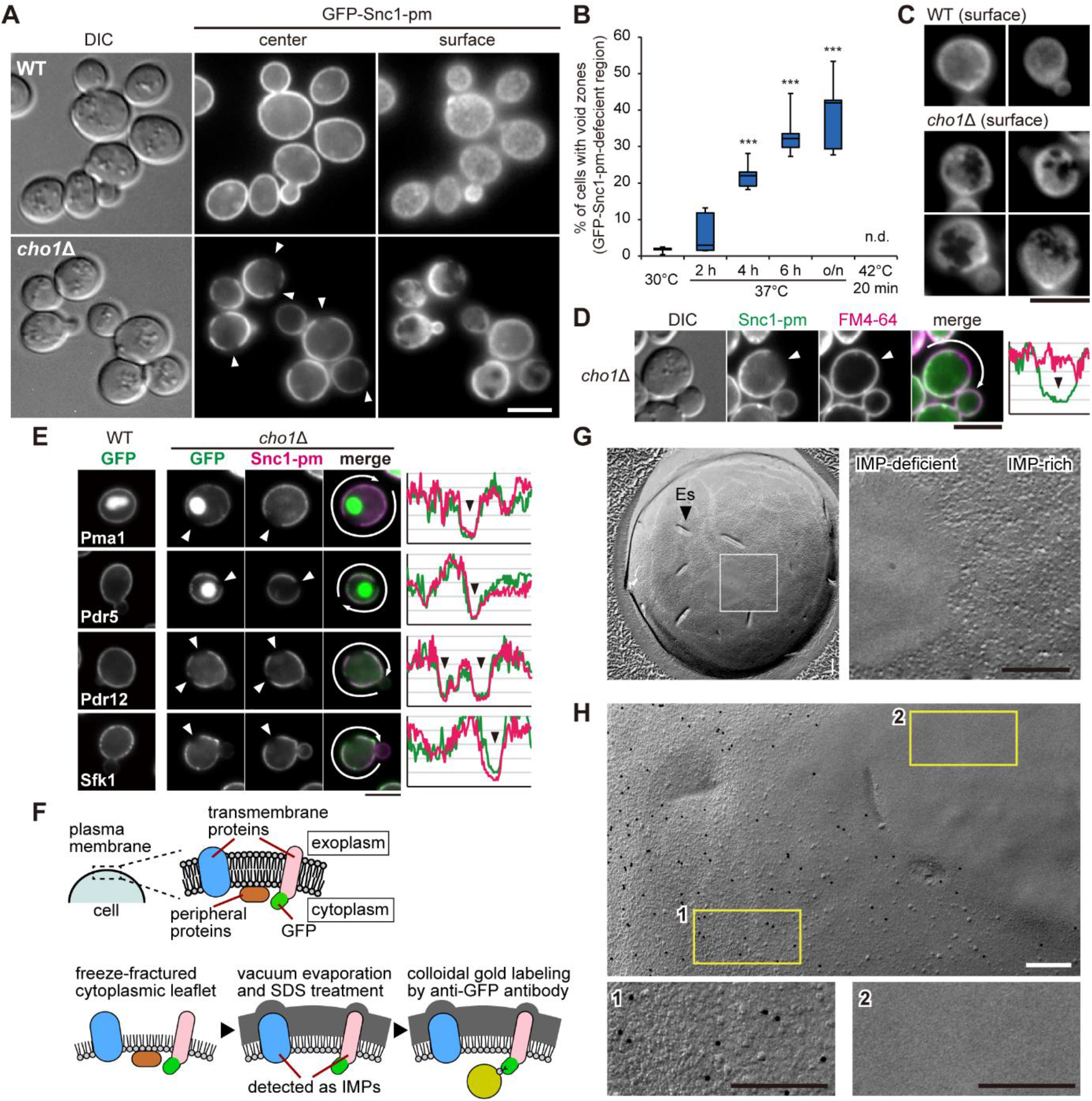
A protein-free region, void zone, is present in the plasma membrane of the PS-deficient yeast cells. (A) Representative images of the void zone. Cells expressing GFP-Snc1-pm were grown overnight in YPDA medium at 37°C. Arrowheads indicate the void zone. Scale bar: 5 μm. (B) The percentage of cells with the void zone. *cho1*Δ cells expressing GFP-Snc1-pm were grown under the indicated conditions to 0.8-1.5 OD_600_. The incidence of the void zone was examined (n > 100 cells, five independent experiments) and is shown as a box plot. Asterisks indicate significant differences from the data obtained at 30°C according to the Tukey–Kramer test (***, p < 0.001; n.d., not detected). (C) Representative images of the void zone in the cell surface. Cells were prepared as in (A). Scale bar: 5 μm. (D) The lipophilic dye FM4-64 can be distributed in the void zone. *cho1*Δ cells were prepared as in (A) and stained with FM4-64 just before the observation. Fluorescence intensities of GFP-Snc1-pm and FM4-64 around the cell (arrows) are plotted on the right. Arrowheads indicate the void zone. Scale bar: 5 μm. (E) The void zone is common to various transmembrane proteins. Cells expressing the indicated proteins were prepared as in (A). Arrowheads indicate the void zone. Fluorescence intensities around the cell (arrows) were plotted as in (D). Scale bar: 5 μm. (F) A scheme of freeze-fracture replica labelling method. (G, H) Freeze-fracture EM images of the plasma membrane of *cho1*Δ cells (G) or Pmal-GFP-expressing *cho1*Δ/*P_GAL1_-CHO1* diploid cells (H). Cells were grown at 37°C. The enlarged image of the area indicated by the square is shown on the right (G) or at the bottom (H). Colloidal gold particles indicate Pma1-GFP labelled by anti-GFP antibody (H). Scale bars: 0.2 μm. (IMP, intramembrane particles; Es, eisosome)

We also investigated the distribution of eisosomes, large immobile protein complexes that form furrow-like invaginations in the fungal plasma membrane (Douglas and Konopka, 2014). Eisosome components, Pil1 and Sur7, were not distributed in the void zone (Figure S1). The void zone was also devoid of the eisosome structure.

To further examine whether transmembrane proteins are completely absent in the void zone, electron microscopy combined with the freeze-fracture replica method was applied (Fujita *et al*., 2010; Tsuji *et al*., 2017). In this method, transmembrane proteins are detected as the granular structures called intramembrane particles (IMPs) (Figure 1F). In the protoplasmic face of the plasma membrane in *cho1*Δ cells, most of the regions were IMP-rich although a submicron-sized smooth region without IMPs was detectable (Figure 1G). To examine whether this IMP-deficient area is the void zone, we labelled Pma1-GFP with an anti-GFP antibody and colloidal gold-conjugated protein A (Figure 1F). As expected, only the IMP-rich area was stained for Pma1-GFP and there was no labelling in the IMP-deficient area (Figure 1H). This result is consistent with the fluorescence microscopy images (Figure 1E) and indicates that the void zone corresponds to the IMP-deficient area. Thus, there are no transmembrane proteins in the void zone.

### Characterization of the dynamics of the void zone

We next examined the stability of the void zone by time-lapse imaging. Most of the void zones were detected over tens of minutes, and some void zones were detectable for longer period of time over 90 min. Bud growth and cytokinesis appeared to proceed normally in the void zone-containing cells, suggesting that the void zone does not influence cell cycle progression (Figure 2A). Consistently, *cho1*Δ cells showed no significant growth defects at 37°C compared to growth at 30°C (Figure 2B). The void zone rarely formed at 30°C (Figure 1B), but once formed, the void zone was largely maintained even after 90 min of incubation at 30°C (Figure. 2C). These results suggest that the void zone is a very stable lipid domain. On the other hand, the void zone was essentially undetectable in a saturated culture with high cell density and very low cell growth. Thus, we investigated whether the frequency of the void zone formation is associated with the growth phase (Figure 2D). Yeast cells show rapid growth in the presence of abundant carbon sources such as glucose. When glucose is depleted, the cells switch to a slower growth rate using ethanol as a metabolic carbon source via a diauxic shift and consequently enter the stationary phase (Gray *et al*., 2004). As shown in Figure 2D, the frequency of the void zone detection was increased early in the logarithmic growth phase and began to decrease starting from the middle phase. Remarkably, the void zone rapidly disappeared when the cells entered the diauxic shift, and no changes were observed after 12 hours, suggesting that the void zone is formed and maintained only in the presence of abundant carbon sources. To investigate the disappearance of the void zone, we examined whether certain stresses induce the disappearance. *cho1*Δ cells grown at 37°C were incubated for additional 3 h at 37°C under three stress conditions, including ATP depletion, glucose starvation, and translation inhibition; then, the cell numbers and the frequency of the void zone were measured. Cell growth was almost completely blocked by all these stresses, and the frequency of the void zone detection was significantly reduced by ATP depletion and glucose starvation, but not by translation inhibition (Figure 2E). These results suggest that the maintenance of the void zone is energy-dependent and does not necessarily require cell growth.

**Figure 2.**
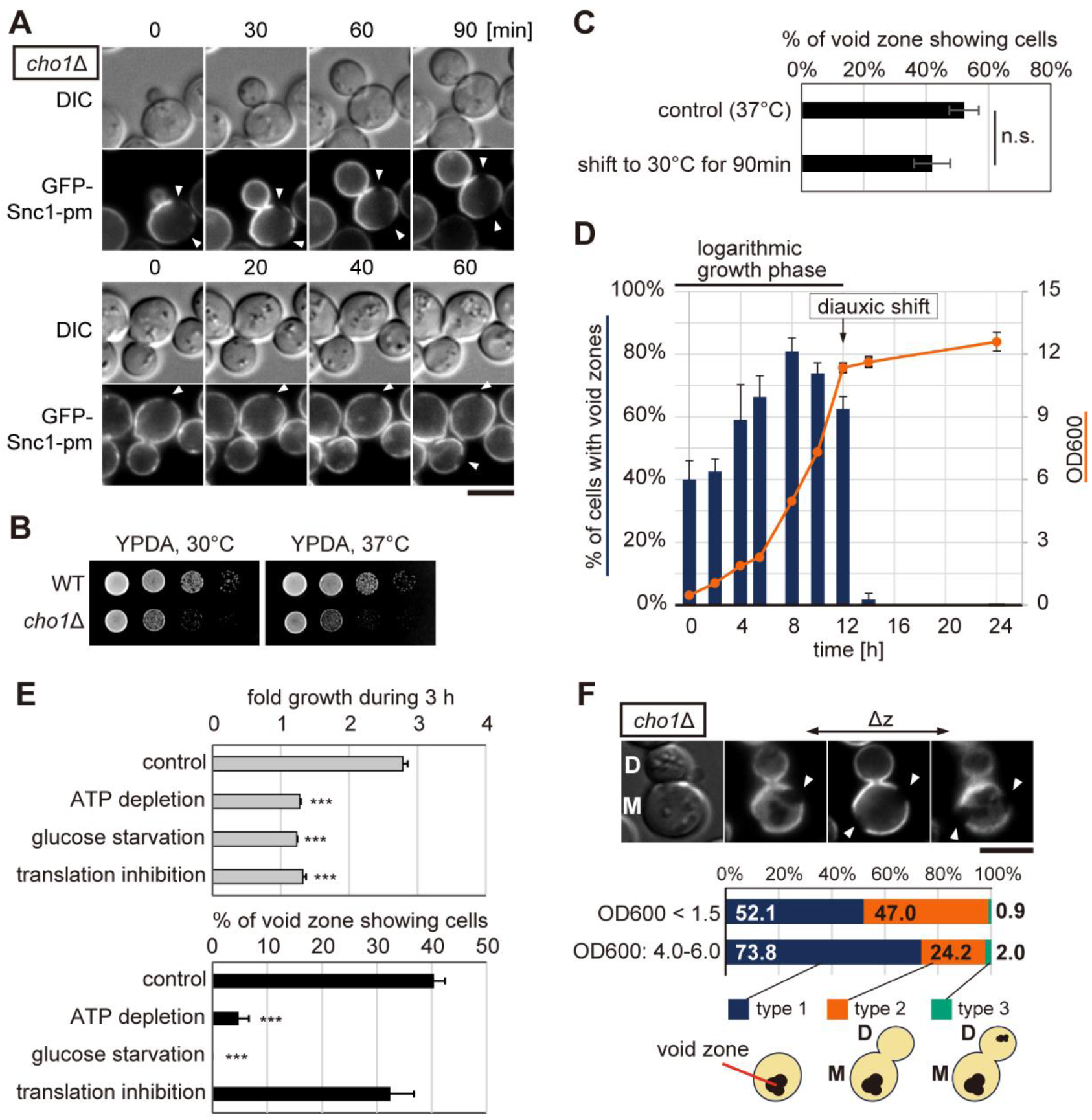
The void zone is stably maintained in an energy-dependent manner. (A) Time-lapse imaging of the void zone. *cho1*Δ cells expressing GFP-Snc1-pm were grown in YPDA medium at 37°C. Two representative examples are shown; a cell with a growing bud (upper panels) and a dividing cell (lower panels) with the void zones. Arrowheads indicate the void zone. Scale bar: 5 μm. (B) Normal growth of *cho1*Δ cells at 37°C. Serial dilutions of cultures were spotted onto the YPDA plates followed by incubation at 30°C or 37°C for 1 d. (C) The void zone formed at 37°C remains stable at 30°C. *cho1*Δ cells expressing GFP-Snc1-pm were grown in YPDA at 37°C, and then shifted to 30°C for 90 min. The percentage of cells with the void zone before and after incubation at 30°C was measured. Data of three independent experiments (n > 100 cells each) are shown as the mean and SD. (D) Frequency of the void zone generation at various growth stages. *cho1*Δ cells expressing GFP-Snc1-pm were grown in YPDA at 37°C, and the percentage of the cells with void zones (blue bars) and the OD_600_ (orange line) were determined. Data of three independent experiments (n > 100 cells each) are shown as the mean and SD. (E) Disappearance of the void zone under stress conditions. *cho1*Δ cells expressing GFP-Snc1-pm were grown to the early log phase at 37°C and were shifted to YPDA medium containing 20 mM sodium azide (ATP depletion), YPA medium lacking glucose (glucose starvation), and YPDA medium containing 10 μg/ml cycloheximide (translation inhibition). After incubation for 3 h at 37°C, OD_600_ and the incidence of the void zone (n > 100 cells) were determined. Data of three independent experiments are shown as the mean and SD. Asterisks indicate significant differences determined by the Tukey–Kramer test (***, p < 0.001). (F) Biased formation of the void zone in the mother cell. *cho1*Δ cells expressing GFP-Snc1-pm were prepared as in (A). A representative pattern of the void zone taken on the different Z focal planes is shown in the upper panel; the void zone occurs in the mother cell (M) but not in the daughter cell (D). Arrowheads indicate the void zone. Scale bar: 5 μm. The percentage of the cells with the void zone in the indicated pattern types (n > 200 cells each) is shown in the lower panel.

We also noticed that the void zone tends to occur in the mother cells rather than the daughter cells (Figure 2F). To confirm this, the distribution of the void zone was categorized into the following groups; cells with no bud or small bud (type 1), cells with mid-large bud in which the void zone occurs only in the mother cell (type 2), and cells in which the void zone occurs in the mother and daughter cells (type 3). As a result, almost all void zones appeared only in the mother cells before or during budding (type 1 and 2); very few type 3 cells were detected, and no cells with void zones formed only in the daughter cells were observed (Figure 2F). This biased distribution was observed even under the condition of OD_600_ = 4.0-6.0, which shows high frequency of the void zone. Exocytosis and endocytosis frequently occur in a bud, which is a polarized growth site, compared to a mother cell. These results suggest that the void zone is generated in a more static membrane.

### Void zone formation is specific to PS-depletion and is a reversible process

To examine whether the void zone can be detected in the lipid mutants other than *cho1*Δ, we investigated the mutants of the genes involved in the synthesis of phosphatidylethanolamine (PE) and phosphatidylcholine (PC). In PE-, and PC-deficient cells, GFP-Snc1-pm was evenly distributed in the plasma membrane, and the void zone was not detected (Figure 3A). We next examined whether the recovery of PS levels dissipates the void zone by adding lyso-PS to the culture media. Incorporated lyso-PS is rapidly converted to PS in the endoplasmic reticulum (ER) via a *CHO1-*independent pathway (Fairn *et al*., 2011; Maeda *et al*., 2013). The recovery of PS was verified by expressing a PS-specific biosensor Lact-C2 (Yeung *et al*., 2008). In *cho1*Δ cells without lyso-PS, the frequency of the void zone did not change for 60 min and mRFP-Lact-C2 remained diffuse in the cytosol. On the other hand, the addition of lyso-PS significantly reduced the incidence of the void zone and resulted in the localization of mRFP-Lact-C2 to the plasma membrane (Figure 3B). These results indicate that the void zone formation is a reversible process. After the addition of lyso-PS, void zones did not disappear uniformly, but rather appeared to be gradually repaired from the boundaries (Figure 3C). In some *cho1*Δ cells, mRFP-Lact-C2 was distributed outside of the void zone (Figure 3D). This void zone is gradually repaired by PS around the void zone. Our results suggest that the void zone is a lipid domain that consists of a lipid (lipids), and random distribution of these lipids in the plasma membrane is facilitated by interaction with PS.

**Figure 3.**
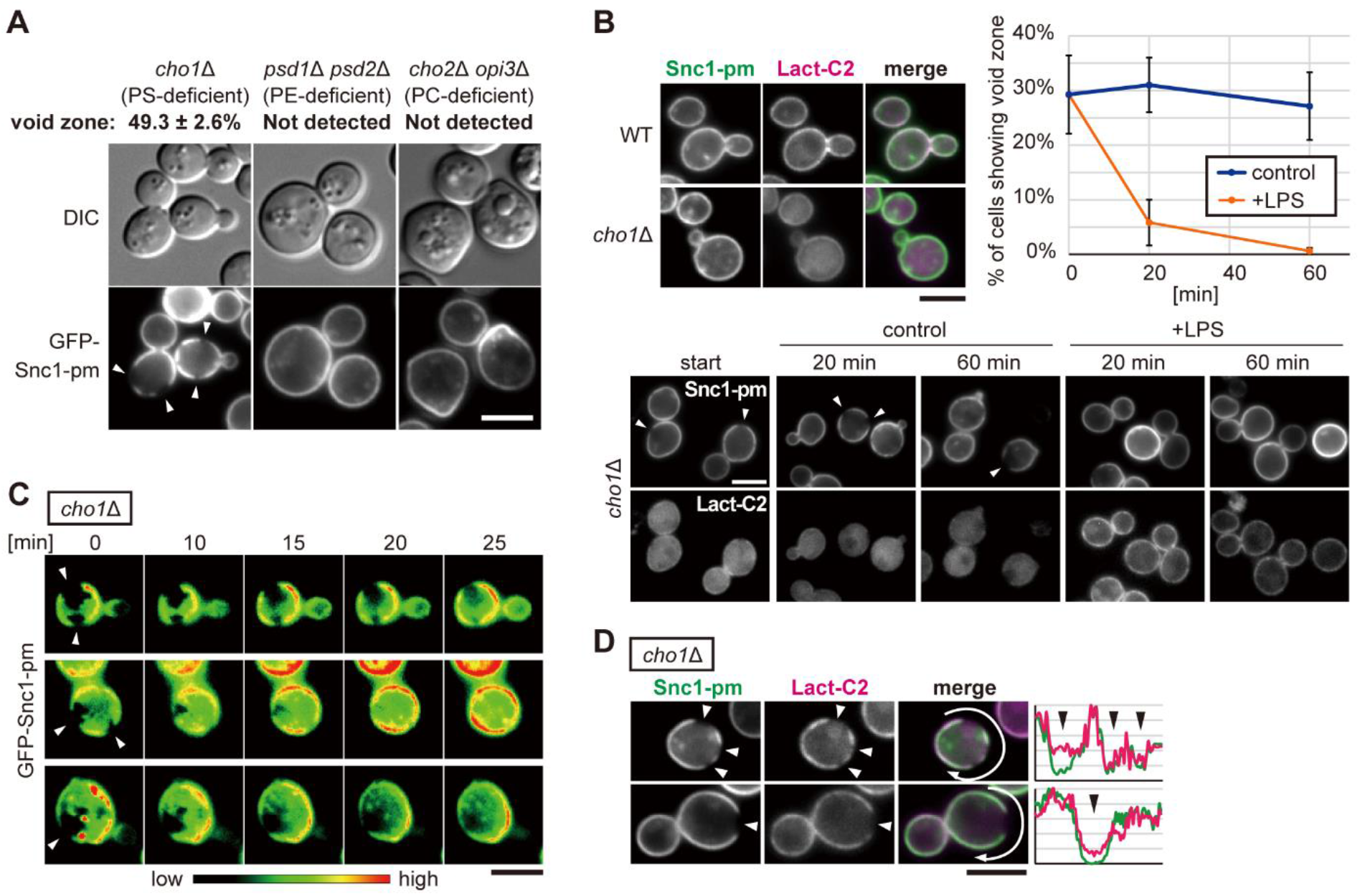
Void zone formation is specific to PS depletion and is a reversible process. (A) The generation of the void zone is specific to the PS-deficient cells. All strains expressing GFP-Snc1-pm were grown at 37°C. *cho1*Δ cells were grown in SD medium containing 1 mM ethanolamine. *psd1*Δ *psd2*Δ cells were grown in SD medium containing 2 mM choline. *cho2*Δ *opi3*Δ cells were grown in SD medium. Percentage of the cells with the void zone is indicated as the mean and SD (n > 100 cells, three independent experiments). Arrowheads indicate the void zone. Scale bar: 5 μm. (B) Regeneration of PS by lyso-PS supplementation dissipates the void zone. Wild-type and *cho1*Δ cells expressing GFP-Snc1-pm and mRFP-Lact-C2 are shown in the upper left panel. *cho1*Δ cells were grown in YPDA medium at 37°C in the presence or absence of lyso-PS (20 μM). Percentage of the cells with the void zone was examined at 20 and 60 min and shown as the mean and SD (n > 100 cells, three independent experiments) (upper right panel). Representative images are shown in the lower panel. Arrowheads indicate the void zone. Scale bars: 5 μm. (C) Time-lapse imaging of the void zone disappearance induced by lyso-PS addition. *cho1*Δ cells expressing GFP-Snc1-pm were prepared as in (B). Cell suspension was placed on the agarose gel pad immediately after mixing with lyso-PS and time-lapse imaging was started. Three examples are shown in the pseudo-colour. Arrowheads indicate the void zone. Scale bar: 5 μm. (D) Void zones are not rapidly dissipated by PS. *cho1*Δ cells expressing GFP-Snc1-pm and mRFP-Lact-C2 were prepared as in (B). Images 20 min after supplementation with lyso-PS are shown. Fluorescence intensities of GFP and mRFP in the cell periphery (arrows) are plotted on the right. Arrowheads indicate the void zone. Scale bar: 5 μm.

### Void zone restricts lateral diffusion of the inner leaflet-anchored proteins, but has no effect on a GPI-anchored protein

We next examined whether the plasma membrane-localized peripheral membrane proteins can be distributed in the void zone. Three types of proteins were tested (Figure 4A). Ras2 is a small GTPase with the C-terminal lipid moiety inserted into the cytoplasmic leaflet of the plasma membrane. Ras2 undergoes farnesylation and palmitoylation, and the latter is required for membrane localization (Bhattacharya *et al*., 1995). Gap1C is the C-terminal cytosolic region of the amino acid permease Gap1. Gap1 localizes to the plasma membrane, and Gap1C without the transmembrane domain is also localized at the plasma membrane (Popov-Čeleketić *et al*., 2016). Gap1C is palmitoylated; however, its membrane localization is due to its amphipathic helix structure and not to the palmitoyl anchor (Popov-Čeleketić *et al*., 2016). Gas1 is a cell wall protein with glycosylphosphatidylinositol (GPI)-anchor inserted into the extracellular leaflet of the plasma membrane (Nuoffer *et al*., 1991). All these proteins were mainly localized to the plasma membrane in the wild-type cells (Figure 4B). In *cho1*Δ cells harbouring the void zone, Ras2 and Gap1C were absent from the void zone similar to Snc1-pm (Figure 4B). In contrast, Gas1 was uniformly distributed in the plasma membrane regardless of the void zone (Figure 4B). These results suggest that the void zone restricts the lateral diffusion of the inner leaflet-associated proteins and the transmembrane proteins but does not influence the diffusion of proteins anchored to the outer leaflet. Normally, PS is abundantly distributed in the inner leaflet of the plasma membrane. These results suggest that formation of the void zone is mainly due to lipid changes in the inner leaflet rather than that in the outer leaflet.

**Figure 4.**
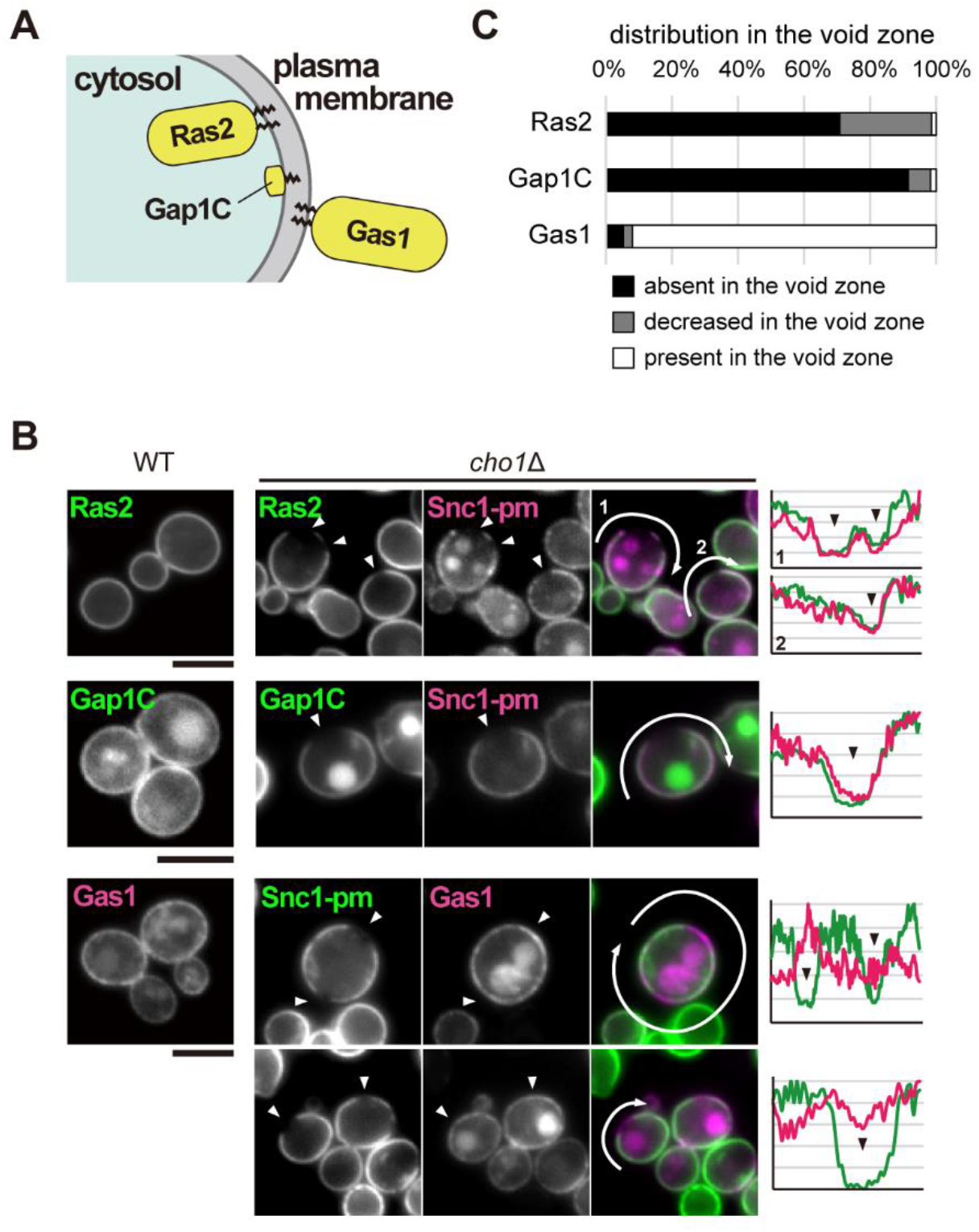
Distribution of peripheral membrane proteins in the void zone. (A) Localization of three peripheral membrane proteins. (B) Ras2 and Gap1C were excluded from the void zone, but Gas1, a GPI-AP, was not excluded. Cells expressing GFP-Ras2 under the control of the *GAL1* promoter and mRFP-Snc1-pm were grown in YPGA at 37°C. Cells expressing GFP-Gap1C and mRFP-Snc1-pm or GFP-Snc1-pm and mRFP-Gas1 were grown in SDA-U medium containing 1 mM ethanolamine at 37°C. Fluorescence intensities of GFP and mRFP in the cell periphery (arrows) are plotted on the right. Arrowheads indicate the void zone. Scale bars: 5 μm. (C) Cells observed as in (B) are categorized by the distribution of each protein in the void zones detected by Snc1-pm (n > 50 cells with the void zone).

### Void zone requires ergosterol and sphingolipid for its formation

Molecular dynamics computer simulation, the giant unilamellar vesicle (GUV) assay, and giant plasma membrane vesicle (GPMV) assay have shown that transmembrane helices and transmembrane proteins are excluded from cholesterol-enriched lipid domains (Sengupta *et al*., 2008; Schäfer *et al*., 2011). Transmembrane proteins are also excluded from ergosterol-enriched raft-like domains that are generated in the vacuolar membrane of yeast in stationary phase (Toulmay and Prinz, 2013; Wang *et al*., 2014; Tsuji *et al*., 2017). Therefore, we examined whether the void zone is a sterol-enriched lipid domain. To assess the distribution of ergosterol, a major sterol in yeast, cells were stained with the sterol-binding dye filipin. Filipin showed a punctate distribution in the plasma membrane of the wild-type cells as reported previously (Grossmann *et al*., 2007), whereas in *cho1*Δ cells, filipin mainly accumulated at the void zones (Figure 5A), indicating that the void zone is rich in ergosterol.

**Figure 5.**
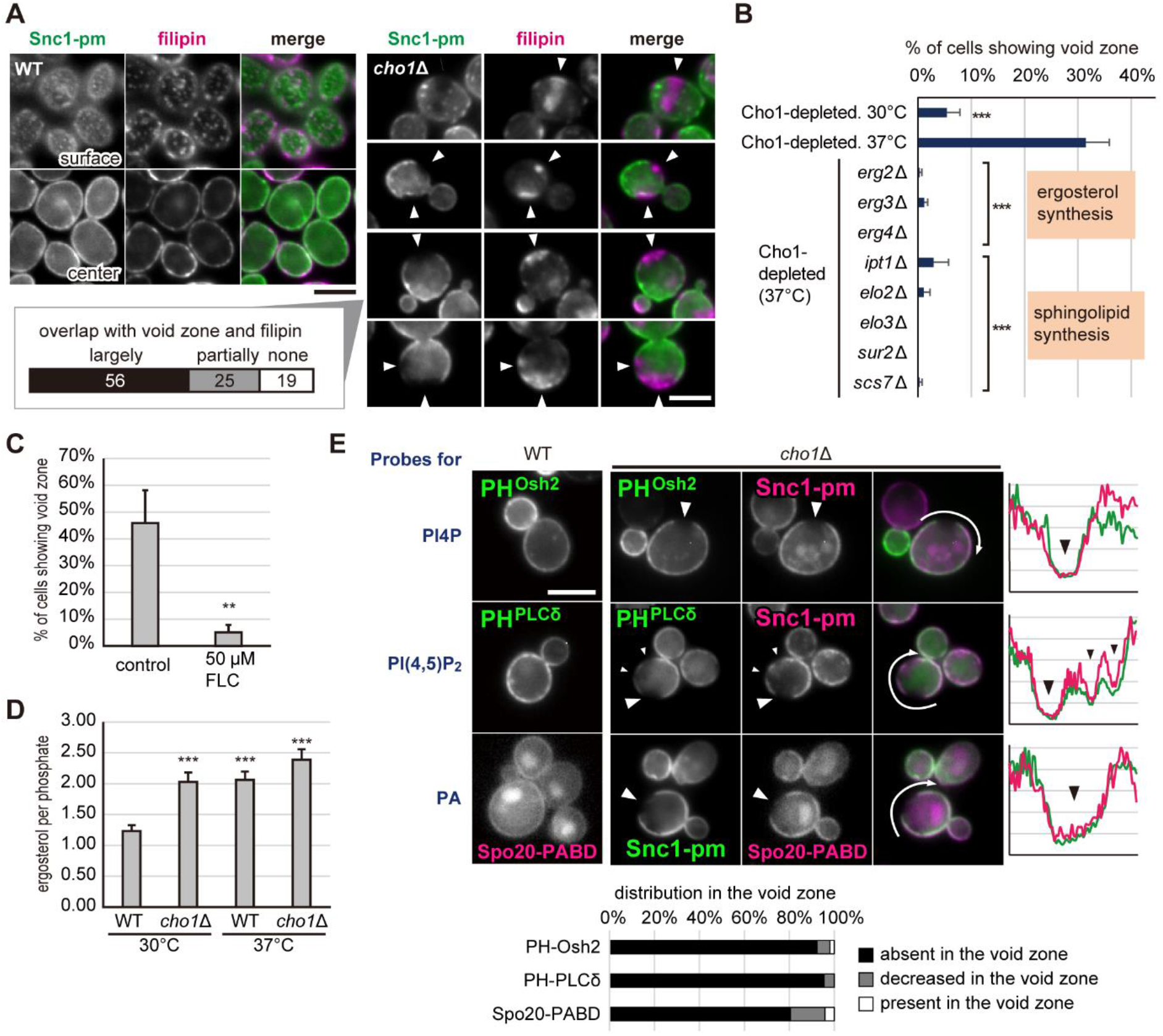
Ergosterol is accumulated in the void zone. (A) Ergosterol accumulation in the void zone. Cells expressing GFP-Snc1-pm were grown in YPDA medium at 37°C, fixed, and stained with filipin. The filipin staining patterns of *cho1*Δ cells containing the void zone (n = 100) were categorized and are shown in the lower left panel. Arrowheads indicate the void zone. Scale bars: 5 μm. (B) The formation of the void zone was dependent on ergosterol and sphingolipids. Cells expressing GFP-Snc1-pm were grown overnight in YPDA medium at the indicated temperature. The incidence of the void zone (n > 100 cells) was examined in three independent experiments, and the data are shown as the mean and SD. Asterisks indicate significant differences from the Cho1-depleted cells grown at 37°C according to the Tukey–Kramer test (***, p < 0.001). (C) The void zone was not generated after ergosterol depletion by fluconazole. *cho1*Δ cells expressing GFP-Snc1-pm were precultured in YPDA medium at 30°C and incubated at 37°C for 6 h with or without 50 μM fluconazole. The incidence of the void zone (n > 100 cells) was examined and the data are shown as in (B). Asterisks indicate a significant difference according to the Student’s t-test (**, p < 0.005). (D) The total levels of ergosterol in the wild-type and *cho1*Δ cells. Cells were grown overnight in YPDA medium at 30°C or 37°C, and total cellular lipids were extracted. Ergosterol contents were analysed by TLC. The ratio of total ergosterol to total phosphate (mol/mol) is shown as the mean and SD of four independent experiments. Asterisks indicate significant differences according to the Tukey–Kramer test (***, p < 0.001 compared to the wild-type at 30°C). (E) PIPs and PA were excluded from the void zone. Cells expressing mRFP-Snc1-pm and either Osh2-2xPH-3xGFP or GFP-PLCδ-2XPH were grown and observed as in (B). Cells expressing GFP-Snc1-pm and mCherry-Spo20-PABD were grown in SDA-U medium containing 1 mM ethanolamine at 37°C. Fluorescence intensities of GFP and either mRFP or mCherry around the cells (arrows) are plotted on the right. Arrowheads indicate the void zone. Scale bar: 5 μm. The distribution of each probe in the void zones detected by Snc1-pm (n > 50 cells with the void zone) is shown in the lower panel.

Similar to sterols, sphingolipids are important for the formation of specialized membrane domains (Dietrich *et al*., 2001; Baumgart *et al*., 2003; de Almeida *et al*., 2003; Veatch and Keller, 2003). We therefore examined whether the biosynthesis of ergosterol and sphingolipid is required for the formation of the void zone. Erg2, Erg3, and Erg4 are essential enzymes for the synthesis of ergosterol (Silve *et al*., 1996; Arthington *et al*., 1991; Lai *et al*., 1994). Ipt1 is an inositolphosphotransferase required for the synthesis of mannosyl-diinositolphosphorylceramide (M(IP)2C), the most abundant sphingolipid in yeast (Dickson *et al*., 1997). Elo2 and Elo3, fatty acid elongases, participate in the long chain fatty acid biosynthesis of sphingolipids (Oh *et al*., 1997). Sur2 and Scs7 are hydroxylases involved in the hydroxylation of sphingolipids (Haak *et al*., 1997). Disruption of any of these genes essentially abolished the formation of the void zone (Figure 5B). Consistently, an inhibitor of ergosterol synthesis, fluconazole, blocked the formation of the void zone (Figure 5C). These results indicate that ergosterol and sphingolipids are necessary for the formation of the void zone and suggest that the void zone may be a novel lipid domain composed of abundant ergosterol and sphingolipids.

We further investigated whether the formation of the void zone results from an increase in the ergosterol level. The ergosterol level was increased in the *cho1*Δ cells grown at 37°C compared to that in the wild-type cells grown at 30°C. However, a similar increase in the ergosterol level was observed in the wild-type cells grown at 37°C and *cho1*Δ cells grown at 30°C, in which the void zone was not formed (Figure 5D). Therefore, the generation of the void zone is not closely correlated with an increase in ergosterol.

The finding that the void zone is an ergosterol-rich lipid domain prompted us to investigate the distribution of other lipids that can be visualized using the corresponding probes. To examine the distribution of phosphatidylinositol 4-phosphate (PI(4)P), phosphatidylinositol 4,5-bisphosphate (PI(4,5)P2), and phosphatidic acid (PA), the PH domains of yeast Osh2 and human PLCδ and the PA-binding domain of yeast Spo20 were used as the probes, respectively (Roy and Levine, 2004; Lemmon *et al*., 1995; Watt *et al*., 2002; Nakanishi *et al*., 2004). These probes were uniformly distributed in the plasma membrane in the wild-type cells; however, none of the probes were detected in the void zone (Figure 5E). This result suggests that PI(4)P, PI(4,5)P2, and PA cannot enter the void zone by lateral diffusion. The majority of yeast phospholipids have mono- or di-unsaturated fatty acids (Schneiter *et al*., 1999; Ejsing *et al*., 2009; Klose *et al*., 2012). In artificial membrane systems, these unsaturated lipids have also been found to be separated from cholesterol-enriched membrane domains (Baumgart *et al*., 2003; Baumgart *et al*., 2007; Risselada and Marrink, 2008; Lingwood and Simons, 2010).

Our results show that the void zone is a lipid domain with an unusual assembly of ergosterol that cooperates with sphingolipids and thereby has novel properties with limited lateral diffusion of the peripheral membrane proteins and certain types of glycerophospholipids.

### Vacuole, a lysosome-like organelle in yeast, contacts the void zone

Observations using the lipid probes suggest phase separation in the plasma membrane; the lipid composition of the void zone is different from that in the other plasma membrane regions. The plasma membrane and the ER form the membrane contact sites (MCSs) via ER-resident tethering proteins that interact with phospholipids of the plasma membrane (Saheki and De Camilli, 2017). To test whether the void zone can influence this ER-PM contact, we examined two ER marker proteins, Hmg1 and Rtn1 (Koning *et al*., 1996; De Craene *et al*., 2006). The cortical ER (cER) was clearly absent at the void zone in the *cho1*Δ cells (Figure 6A; 89.2% of Hmg1, 92.9% of Rtn1, n > 100 cells, respectively). The ER-PM tethering proteins Tcb1/2/3, and Ist2 were reported to bind to PS and PI(4,5)P2, respectively (Schulz and Creutz, 2004; Fischer *et al*., 2009). Since PS is not synthesized in the *cho1*Δ cells and PI(4,5)P2 is not distributed in the void zone (Figure 5E), the association of cER with the plasma membrane may be lost in the void zone.

**Figure 6.**
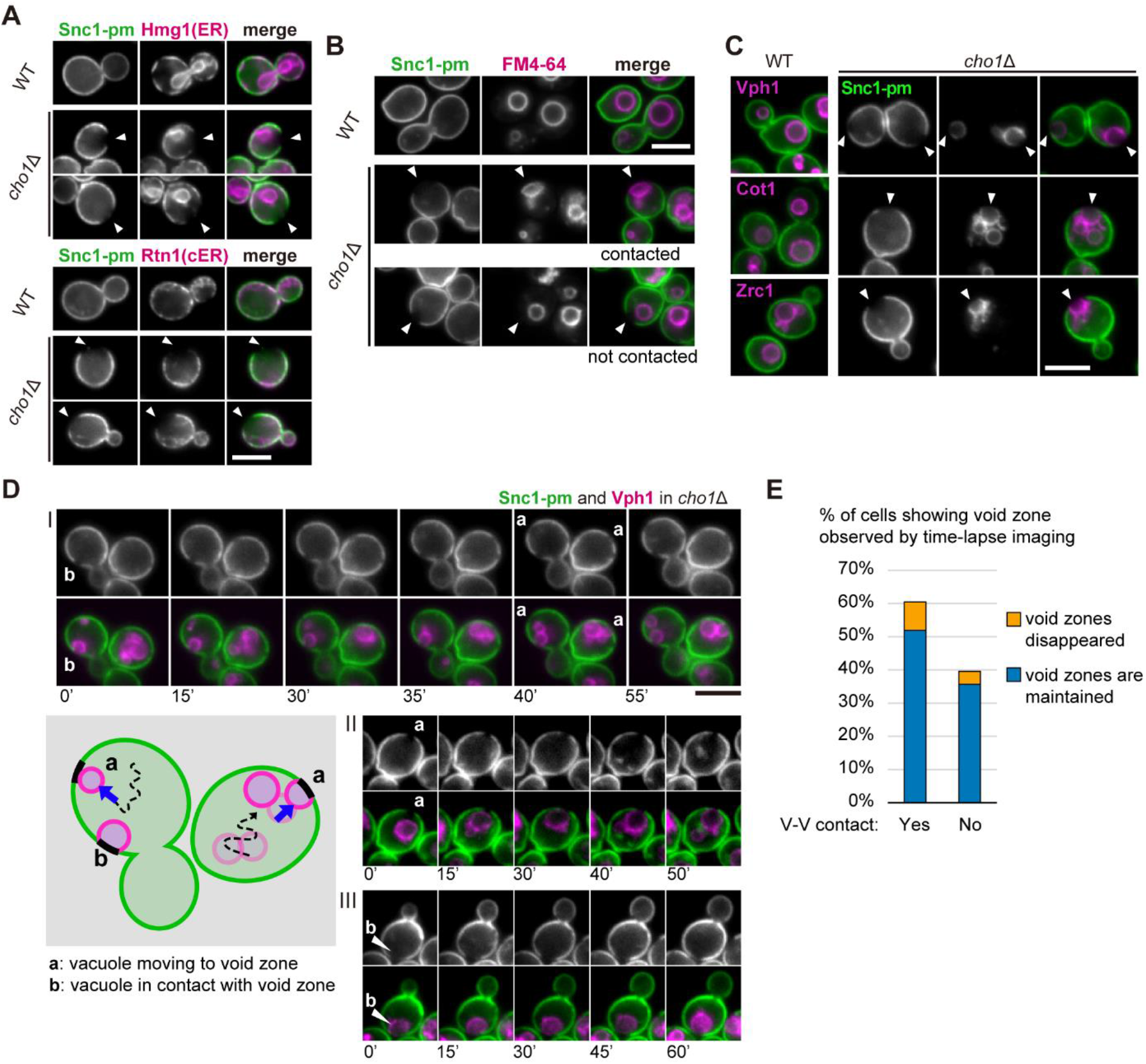
Vacuoles contact the void zone. (A) Cortical ER is disassociated from the void zone. Cells expressing mRFP-Snc1-pm and either Hmg1-GFP or Rtn1-GFP were grown in YPDA medium at 37°C. The mRFP-Snc1-pm and ER marker proteins are shown in green and magenta, respectively. Arrowheads indicate the void zone. Scale bar: 5 μm. (B) Vacuoles contact the void zone. Cells expressing GFP-Snc1-pm were grown in YPDA medium at 37°C and stained with FM4-64 for 20 min. Arrowheads indicate the void zone. Scale bar: 5 μm. (C) Separation of the vacuolar protein in the void zone contact region. Cells expressing mRFP-Snc1-pm and either Vph1-GFP or GFP-Cot1 and cells expressing GFP-Snc1-pm and mRFP-Zrc1 were grown in YPDA medium at 37°C. Snc1-pm and vacuolar proteins are shown in green and magenta, respectively. Arrowheads indicate the void zone. Scale bar: 5 μm. (D) Time-lapse imaging of the vacuole-void zone contact. *cho1*Δ cells expressing GFP-Snc1-pm and Vph1-mRFP were grown in YPDA medium at 37°C. Three examples with two distinctive patterns (a, b) are shown. In the bottom left scheme, black regions in the plasma membrane indicate the void zone. Numbers indicate time in min. Scale bar: 5 μm. (E) The vacuole-void zone contact is stable. Cells observed by time-lapse imaging as in (D) are categorized by the presence or absence of the V-V contact and the void zone behaviour (n > 100 cells with the void zone). Estimation was based on the void zone or the V-V contact lasting more than 30 min during 1 h observation.

To investigate whether the void zone influences the distribution or morphology of other organelles, several organelle markers were examined in the *cho1*Δ cells. Surprisingly, we found that vacuoles were in contact with the void zones (Figure 6B). The observations using Snc1-pm and FM4-64 indicate that the contact between the void zone and the vacuole was detected in 45.1% of the *cho1*Δ cells with the void zone (45.1 ± 7.0%, five independent experiments, n > 50 cells each). The proximity of the non-void zone regions and vacuoles was observed only in 7.8% of the *cho1*Δ cells (7.8 ± 1.7%, three independent experiments, n > 100 cells each) compared to that in the wild-type cells (10.1 ± 2.1%, three independent experiments, n > 100 cells each). These data suggest that frequent contact of the vacuoles with the plasma membrane is specific to the void zone. The fact that not all void zones are in contact with the vacuoles indicates that the exclusion of the transmembrane proteins from the void zones was not caused by the vacuole contact (Figure 6B). On the other hand, the *trans*-Golgi network (TGN), lipid droplets, and mitochondria did not contact the void zone (Figure S2), suggesting that only vacuoles interact with the void zone. This type of contact between the plasma membrane and the vacuoles or lysosomes has not been reported, and we refer to the contact between the void zone and vacuoles as the “V-V contact”.

In stationary phase yeast cells, a raft-like domain is formed in the vacuolar membrane, where lipophagy, the uptake of the lipid droplets, occurs (Toulmay and Prinz, 2013; Wang *et al*., 2014; Tsuji *et al*., 2017). The properties of this vacuolar microdomain are very similar to the properties of the void zone, including absence of the transmembrane proteins and ergosterol enrichment. To test whether this vacuolar microdomain is present in the void zone contact area, we examined three vacuolar transmembrane proteins, Vph1, Cot1, and Zrc1 (Manolson *et al*., 1992; Li and Kaplan, 1998). These proteins were uniformly distributed in the vacuolar membrane of the wild-type cells; however, in some vacuoles in contact with the void zone of the *cho1*Δ cells, these proteins were excluded from the contact area (Figure 6C). One of these proteins, Vph1, had the highest frequency of segregation on vacuoles in contact with the void zone (73.5% of Vph1, 41.5% of Cot1, and 30.0% of Zrc1, n > 50). In contrast, the exclusion of FM4-64 dye in vacuoles of the cells with the V-V contacts was not detected (n > 50; Figure 6B). These data suggest that certain vacuoles form a membrane domain at the V-V contact site.

To understand the dynamics of the V-V contact, we used time-lapse imaging with GFP-Snc1-pm and Vph1-mRFP (Movie 1) and detected two patterns. First, the movement of a vacuole into the void zone was observed (Figure 6D, shown as “a”). The speed of the vacuole migration to the void zone was highly variable between individual cells. However, some void zones had no contact with the vacuoles during the 1 h observation. Second, the V-V contact lasted over half an hour (Figure 6D, shown as “b”). In one hour of observation of the cells with the void zone, 60% of the cells had the V-V contact over 30 min (Figure 6E). The disappearance of the void zone after its formation was rarely observed, and this phenomenon appears to be unrelated to the presence or absence of the V-V contacts. In addition, no vacuoles left the void zone while the void zone was maintained. The molecular basis of the V-V contact is unknown although this contact appears to be stable. Vacuoles in contact with the void zone underwent fission and fusion (Figure S3).

### Identification of the genes required for the formation of the void zone

To further understand the mechanism of the void zone formation, we created a series of deletion mutants on the background of the *cho1*Δ or glucose-repressible *P_GAL1_-3HA-CHO1* mutations and examined their effect on the generation of the void zone (Figure 7A). Various genes involved in sterol trafficking were tested, and the results indicate that *kes1*Δ (*osh4*Δ) mildly and *arv1*Δ significantly reduced the void zone formation. Kes1 is one of the yeast oxysterol-binding proteins that exchanges sterols for PI(4)P between the lipid membranes (Jiang, *et al*., 1994; de Saint-Jean *et al*., 2011). Arv1 was implicated in the GPI-anchor biosynthesis and transport and in intracellular sterol distribution (Kajiwara *et al*., 2008; Beh and Rine, 2004). Both Kes1 and Arv1 are involved in the sterol transport; however, their contribution to the sterol transport to the plasma membrane is very low (Georgiev *et al*., 2011; Georgiev *et al*., 2013). These proteins regulate sterol organization in the plasma membrane; the mutations influence the sensitivity to the sterol-binding drugs and sterol-extraction efficiency of MβCD (Georgiev *et al*., 2011; Georgiev *et al*., 2013). These differences in sterol organization may influence the formation of the void zone. Npc2 is an orthologue of Niemann-Pick type C protein and plays an essential role together with Ncr1 in sterol insertion into the vacuolar membrane from the inside of the vacuole, which is required for the formation of the raft-like vacuolar domain during lipophagy in the stationary phase (Tsuji *et al*., 2017). However, deletion of Npc2 did not influence generation of the void zone or formation of the V-V contacts accompanied with protein-free vacuolar domain (Figure S4). This result suggests that the void zone is formed independently of Ncr1/Npc2-mediated sterol transport possibly because Ncr1/Npc2 are not involved in sterol organization in the plasma membrane. The vacuolar domains detected at the V-V contact region and generated during the stationary phase lipophagy are similar in appearance; however, the processes of their formation may be different.

**Figure 7.**
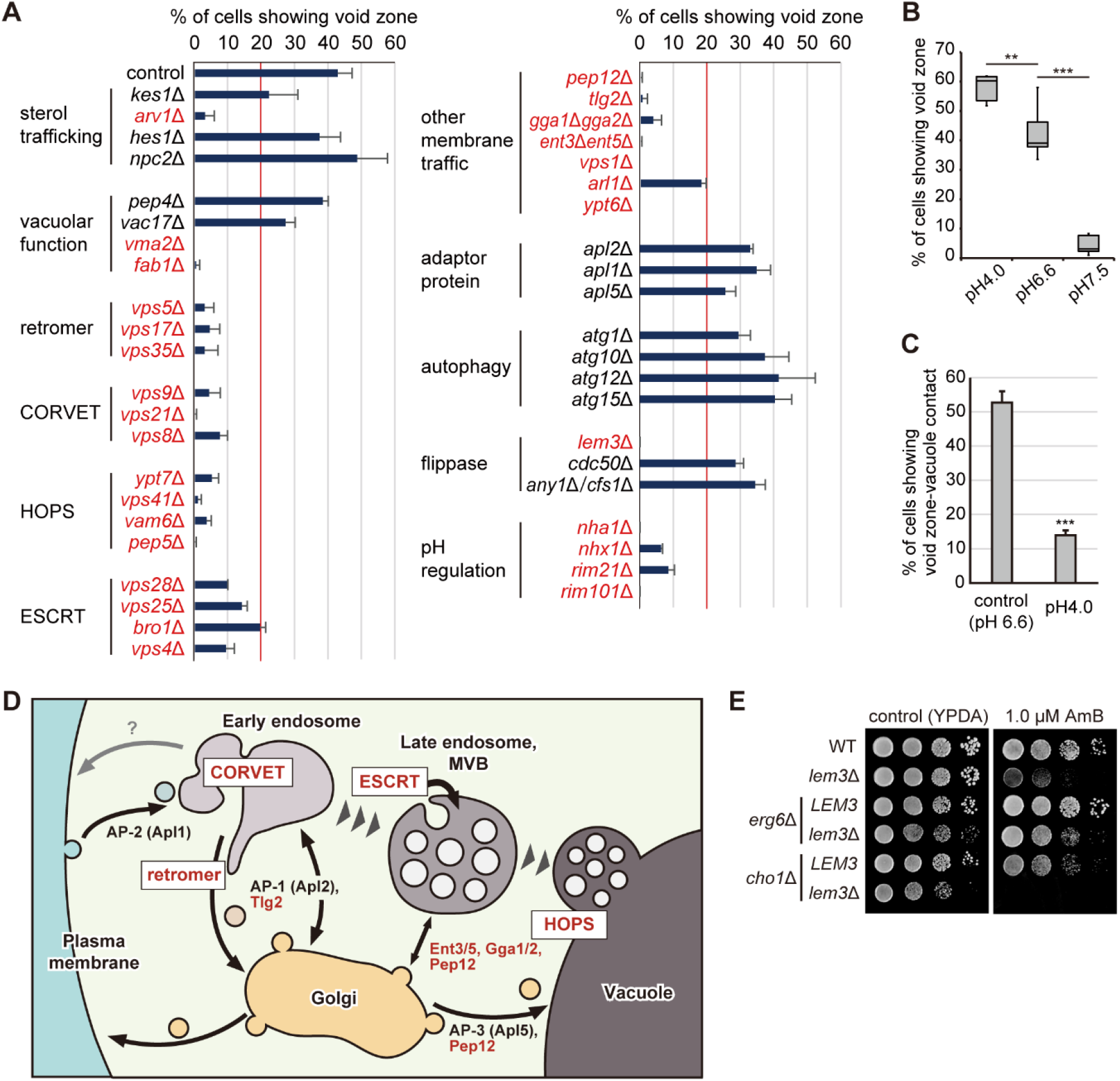
A search for the genes required for the void zone formation. (A) Frequency of the void zone in the mutant cells generated on *cho1*Δ background or under Cho1-depleted conditions. The incidence of the void zone was determined (n > 100 cells, three independent experiments) and is shown as the mean and SD. The mutations responsible for the low incidence (under 20%) are shown in red. (B) The void zone formation is suppressed by high pH. *cho1*Δ cells expressing GFP-Snc1-pm were grown overnight in YPDA at the indicated pH at 37°C. The incidence of the void zone was determined (n > 100 cells, five independent experiments) and is shown as a box plot. Asterisks indicate significant differences according to the Tukey–Kramer test (**, p < 0.005; ***, p < 0.001). (C) Low pH decreases the frequency of the V-V contact. *cho1*Δ cells expressing GFP-Snc1-pm and Vph1-mRFP were grown in YPDA at the indicated pH at 37°C. The incidence of the V-V contacts was determined (n > 100 cells with the void zone, three independent experiments) and is shown as the mean and SD. Asterisks indicate a significant difference according to the Student’s t-test (***, p < 0.001). (D) A scheme of the membrane trafficking. Proteins and protein complexes required for the formation of the void zone are shown in red. (E) The loss of phospholipid asymmetry in the plasma membrane results in high sensitivity to amphotericin B. Serial dilutions of the cultures were spotted onto YPDA plates containing 1.0 μM amphotericin B and incubated at 30°C for 2 d.

The void zone formation was observed in the absence of Pep4, a major vacuolar protease, and Vac17, a myosin adapter required for inheritance of vacuoles. However, the void zone formation was strongly suppressed by deletion of Vma2 and Fab1.

Vma2 is a subunit of the V-ATPase that regulates pH homeostasis and functions as a pH sensor (Marshansky *et al*., 2014). Consistent with this observation, other genes involved in pH homeostasis (*NHA1, NHX1, RIM21*, and *RIM101*) were required for the void zone formation (Figure 7A, pH regulation) (Sychrová *et al*., 1999; Brett *et al*., 2005; Obara *et al*., 2012). Thus, we examined the effect of pH of the medium on the formation of the void zone. The formation of the void zone was strongly suppressed by increasing pH in the medium from 6.6 to 7.5. (Figure 7B). On the other hand, the low pH medium (pH 4.0) slightly increased the formation of the void zone. Interestingly, the frequency of the V-V contacts was significantly reduced in the low pH medium (Figure 7C). The mechanism of these pH-dependent phenomena is unclear although they may be important in assessment of the molecular basis of the formation of the void zones and V-V contacts.

Fab1 is a phosphatidylinositol 3-phosphate (PI(3)P) 5-kinase that generates phosphatidylinositol 3,5-bisphosphate (PI(3,5)P2) (Cooke *et al*., 1998). PI(3,5)P2 functions as a signal lipid in intracellular homeostasis, adaptation, and retrograde membrane trafficking (Jin *et al*., 2016). We speculated that the defects in the retrograde transport may indirectly influence the void zone formation in the plasma membrane and thus examined various genes involved in the membrane transport. Strikingly, conserved protein complexes, retromer, class C core vacuole/endosome tethering (CORVET), homotypic fusion and vacuole protein sorting (HOPS), and endosomal sorting complexes required for transport (ESCRT) were required for the void zone formation (Figure 7A and D). As the name implies, the Vps proteins belonging to these complexes were identified using mutants defective in vacuolar protein sorting (VPS) (Robinson *et al*., 1988; Rothman *et al*., 1989). Dysfunction of these complexes perturbs intracellular vesicle trafficking (Schmidt and Teis, 2012; Balderhaar and Ungermann, 2013; Burd and Cullen, 2014), which may influence the plasma membrane recycling of cargo and lipids involved in the void zone formation. Similarly, proteins involved in the retrograde transport, such as the SNAREs Pep12 and Tlg2, the epsin-like adapter Ent3/Ent5, and the clathrin adapter Gga1/Gga2, were required for the void zone formation. A dynamin-like GTPase Vps1, Arf-like GTPase Arl1, and Rab6 GTPase homologue Ypt6 are known to be closely related to the membrane trafficking (Vater *et al*., 1992; Li and Warner, 1996; Rosenwald *et al*., 2002). Consistent with this notion, *vps1*Δ, *arl1*Δ, and *ypt6*Δ inhibited the void zone formation. Impaired membrane transport in the inner membrane system can be manifested as the changes in the plasma membrane lipid organization and/or defects in the pH control. However, Apl2 and Apl1, the subunits of the adaptor complexes AP-1 and AP-2, respectively, had little contribution to the void zone formation presumably because these mutants had insignificant disruption of the membrane trafficking compared to the effects of *ent3*Δ, *ent5*Δ and *gga1*Δ *gga2*Δ (Yeung *et al*., 1999; Sakane *et al*., 2006; Morvan *et al*., 2015). Deletion of *APL5*, which encodes the subunit of AP-3 responsible for the transport from the Golgi to the vacuole (Dell’Angelica, 2009), slightly reduced the void zone formation, suggesting the importance of the vacuolar functions for the void zone formation.

Deletion of the autophagy-related genes (*atg1*Δ, *atg10*Δ, *atg12*Δ, and *atg15*Δ) did not influence the void zone formation. *ATG1* is one of the core ATG genes (Mizushima *et al*., 2011). The void zone was generated and the V-V contact with the vacuolar microdomain was observed in the absence of Atg1 (Figure 7A, and Figure S4). This result suggests that the void zone formation is independent of autophagy consistent with our notion that direct or indirect effects on lipid organization in the plasma membrane (e.g., *via* the Vps pathway) are influencing the void zone formation.

We also examined the effect of mutations in the flippase-related proteins. The deficiency of Cdc50 or Any1/Cfs1 localized in endosomes and the TGN had little effect on the void zone formation (Saito *et al*., 2004; van Leeuwen *et al*., 2016; Yamamoto *et al*., 2017); however, disruption of Lem3 localized in the plasma membrane completely suppressed the generation of the void zone (Kato *et al*., 2002). The Lem3-Dnf1/2 flippase complexes translocate glycerophospholipids, but not ceramides and sphingolipids, to the cytoplasmic leaflet of the lipid bilayer (Pomorski *et al*., 2003; Saito *et al*., 2004; Furuta *et al*., 2007). We assumed that disruption of the phospholipid asymmetry by *lem3*Δ may influence ergosterol behaviour in the plasma membrane. To test this hypothesis, we examined the sensitivity to an antifungal ergosterol-binding drug, amphotericin B (AmB) (Kamiński, 2014). The results indicate that *lem3*Δ is highly sensitive to AmB, which is detected in the *cho1*Δ background cells, and this effect was cancelled by addition of *erg6*Δ that causes defects in ergosterol biosynthesis (Figure 7E). Thus, the disruption of phospholipid asymmetry alters the ergosterol distribution in the plasma membrane, thereby suppressing the void zone formation (see discussion).

Although it is not yet clear about the mechanisms that some of these additive mutations inhibit the void zone formation, these findings may help to elucidate the nature of the void zone in the future.

## Discussion

In the artificial membranes, phase separation causes micron-scale separation of proteins and lipids; such large separation does not occur in the plasma membrane of the living cells. The mechanism that enables random distribution of proteins and lipids throughout the plasma membranes on the macroscopic scale by preventing large-scale phase separation has not been understood. We found that in PS-deficient cells grown at high temperatures, the protein-free membrane domain “void zone” develops in the plasma membrane and exhibits properties similar to phase-separated artificial membranes (Figure 8). We propose a new role of PS in membrane organization and suggest that PS prevents the occurrence of abnormal membrane domains due to phase separation and ensures macroscopic homogeneity of the distribution of the molecules on the plasma membrane.

**Figure 8.**
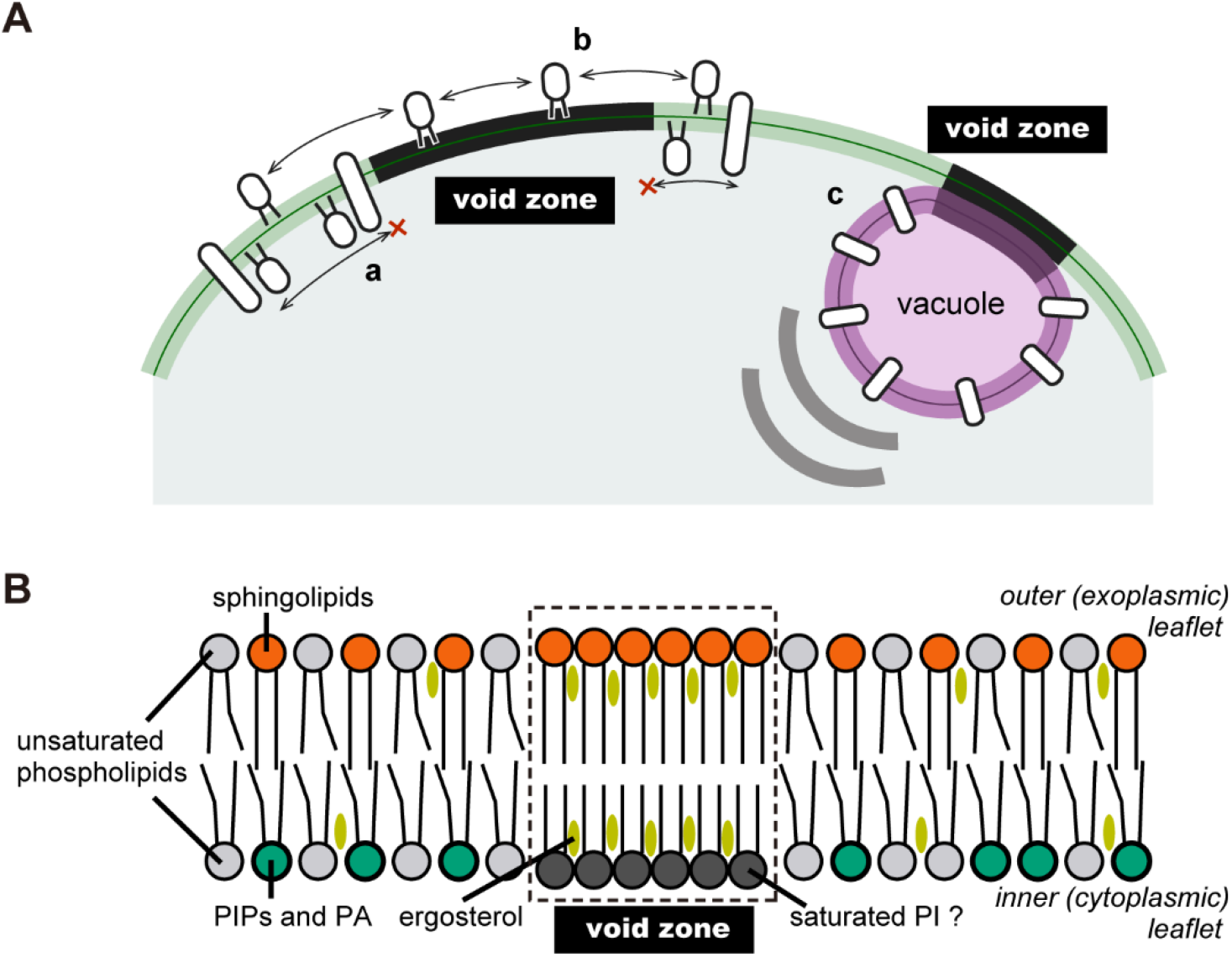
Summary and a model of the void zone. (A) Characteristics of the void zone revealed in this study. Our results suggest the following properties of the void zone: (a) transmembrane proteins and peripheral membrane protein distributed in the inner leaflet cannot enter the void zone by lateral diffusion; (b) a GPI-AP, Gas1, localized in the outer layer of the plasma membrane is not influenced by the void zone; and (c) vacuoles move to form a stable contact with the void zone, and a vacuolar membrane domain also appears to be formed at this contact site. (B) The putative structure of the void zone (see discussion).

### Mechanisms of the generation and disappearance of the void zone

PS is a phospholipid mainly distributed in the inner leaflet of the plasma membrane and has a relatively high affinity for cholesterol (Maekawa and Fairn, 2015; Nyholm *et al*., 2019). Therefore, loss of PS may alter relative affinity of the phospholipids for ergosterol in the inner leaflet and may also influence the transbilayer distribution of ergosterol. Almost all yeast phospholipids, including PS, have at least one unsaturated fatty acid; however, a small fraction of phosphatidylinositol (PI) has only saturated fatty acids (Schneiter *et al*., 1999; Ejsing *et al*., 2009; Klose *et al*., 2012). Molecular species of PS, PE, PC, and PI were characterized in the isolated plasma membrane; interestingly, 29.2% of PI have two saturated fatty acids (Schneiter *et al*., 1999). Relative affinity of ergosterol to various phospholipid species is unclear. One hypothesis is that ergosterol is not clustered due to the interactions with unsaturated PS in the wild-type cells; however, in *cho1*Δ cells, saturated PI becomes the most dominant interaction partner of ergosterol in the inner leaflet of the plasma membrane thus creating a driving force for phase separation. When PS is resynthesized in the *cho1*Δ cells after the addition of lyso-PS, the void zone may disappear because PS becomes the predominant interaction partner of ergosterol (Figure 3B and C).

Generation of the void zone requires PS deficiency and high temperature conditions. In artificial membrane systems, phase separation occurs at a temperature below the miscibility transition temperature, which is dependent on the lipid composition of the membrane (Veatch and Keller. 2002; Baumgart *et al*., 2007; Levental et al., 2009). Contrary to these phenomena, the void zone is rarely formed at 30°C and occurs by incubation at 37°C (Figure 1B); thus, we speculate that a change in miscibility transition temperature due to remodeling of lipid composition at 37°C may be one of the triggers for void zone formation in *cho1*Δ cells. In yeast, high temperature reduces the degree of unsaturation and increases the acyl chain length of glycerophospholipids (Klose *et al*., 2012), and these events promote phase separation in the liposomes (Engberg *et al*., 2016). In addition, our results that several hours of incubation at 37°C was required to generate the void zone (Figure 1B), and that the developed void zone was maintained for 90 min after the temperature was returned to 30°C (Figure 2C), are consistent with this idea and suggest that remodeling of the lipid composition takes a long time. A combination of PS deficiency and high temperature-induced lipid remodelling may create a specific membrane environment that results in the void zone development.

The void zones rapidly disappeared when the *cho1*Δ cells undergo a diauxic shift (Figure 2D) or when glucose or ATP is depleted from the medium (Figure 2E). During glucose starvation, activities of the plasma membrane H^+^-ATPase Pma1 and V-ATPase are reduced (Young *et al*., 2010; Dechant *et al*., 2010). Therefore, the disappearance of the void zone may be caused by a disturbance of pH homeostasis consistent with the suppression of the void zone formation by mutations in the genes involved in pH homeostasis, including *VMA2* (Figure 7A). Reduced activities of Pma1 and V-ATPases lead to acidification of the cytosol, and the void zone formation is also inhibited in the medium with elevated pH (Figure 7B). The relationship between pH and phase separation is largely unknown; however, it has been reported that phase separation occurs at low pH and not at high pH in the artificial membranes that mimic human stratum corneum (Plasencia *et al*., 2007).

Our results indicate that long chain fatty acids and hydroxylation of sphingolipids are necessary for the void zone formation (Figure 5B). It has been reported that GUVs prepared from the yeast total lipids have extensive phase separation, which depends on long fatty acid elongation and hydroxylation of sphingolipids (Klose *et al*., 2010). M(IP)2C with a C26 fatty acid is the most common yeast sphingolipid species and its levels are significantly reduced in the *elo2*Δ and *elo3*Δ mutants (Oh *et al*., 1997; Ejsing *et al*., 2009). A recent study using asymmetric GUVs suggested that C24 SM and not C16 SM has a role in cholesterol retention in the inner leaflet of the lipid bilayer (Courtney *et al*., 2018). Consistently, dehydroergosterol, a closely related fluorescent analogue of ergosterol, is mainly located in the inner leaflet of the plasma membrane, and this asymmetry is maintained by sphingolipids in yeast (Solanko *et al*., 2018). This type of interaction of ergosterol with sphingolipids may be necessary for the void zone formation.

Analysis using several mutant strains on the *cho1* background revealed that retrograde intracellular trafficking has a significant effect on the void zone formation (Figure 7A). Perturbed membrane transport may alter the distribution of the proteins and the key lipids as described above. Moreover, *ARV1* and *LEM3* are important for the void zone formation. In *arv1*Δ cells and Δtether cells that lack the ER-PM contacts, the transport of ergosterol to the plasma membrane is essentially unchanged, whereas there were significant changes in ergosterol accessibility to the compounds, such as increased efficiency of ergosterol extraction with MβCD and increased sensitivity to the sterol-binding drug nystatin (Georgiev *et al*. 2013; Quon *et al*., 2018). Similarly, *lem3*Δ does not change the total ergosterol level (Mioka *et al*., 2018), and Dnf1 and Dnf2 do not contribute to the asymmetric distribution of the ergosterol analogues across the plasma membrane bilayer (Solanko *et al*., 2018). However, *lem3*Δ cells are highly sensitive to AmB (Figure 7E). Thus, similar to the *arv1*Δ and Δtether mutations, *lem3*Δ may alter the ergosterol distribution in the plasma membrane thus preventing the formation of the void zone.

### Is the void zone a liquid-ordered domain?

To date, there are no known examples of micron-scale phase separation in the plasma membrane of living cells, and its reason is not clear (Lee et al., 2015; Levental et al., 2020). Thus, the void zone may be the first report of a micron-scale phase separation in the plasma membrane in vivo. The following features of the void zone are similar to those of the liquid-ordered domain in phase-separated artificial membranes: 1) absence of the transmembrane proteins, 2) absence of certain peripheral membrane proteins, 3) exclusion of certain types of glycerophospholipids, and 4) enrichment in sterols and the contribution of sphingolipids (Baumgart *et al*., 2007; Sengupta *et al*., 2008; Kaiser *et al*., 2009; Schäfer *et al*., 2011). On the other hand, the formation of void zones requires lipid asymmetry and their maintenance is energy-dependent (Figure 2E and 7A), which is a very different feature from the membrane domains in artificial membrane systems. Differences between living and artificial membranes regarding phase separation have been reported previously; macroscopic phase separation does not occur in living wild-type yeast cells (Figure 1A and C), but occurs in GUVs formed with lipids extracted from the wild-type strains (Klose et al., 2010). Similarly, whereas no phase separation is observed in living mammalian cells over a wide range of temperatures, GPMVs prepared from the same cells shows macroscopic phase separation (Lee et al., 2015). Further studies of the void zone would help to bridge the gap between the artificial and biological membranes. Besides the PS-sterol interaction, there may be other underlying mechanisms by which living cells avoid macroscopic phase separation.

### Membrane contact between the void zone and vacuoles

We have detected the contact between the void zone and the vacuoles (Figure 6B and D, Figure 8A), and the time-lapse imaging revealed that this contact lasted for at least 30 min in most cases (Figure 6D and E). These results suggest that the contact vacuoles may not contribute to the rapid degradation and disappearance of the void zone. The contact vacuoles are more likely to play a role in sealing the void zone to protect the cells. Recent studies reported that in the phase-separated membranes with the liquid-disordered and liquid-ordered domains, membrane permeability is increased at the interface between the two domains (Cordeiro, 2018). A similar result indicated that the ER-PM contact sites are important for the maintenance of the integrity of the plasma membrane (Omnus *et al*., 2016; Collado *et al*., 2019). In this study, we found that void zone has Lo phase-like properties and that cortical ER is dissociated from the void zone. Therefore, it is possible that the void zone induces high local permeability on its borders and low local integrity; thus, the void zone may be a fragile region of the plasma membrane. The V-V contact may represent a protective cellular response to these plasma membrane abnormalities.

Our results also suggest that membrane domains may be formed on the vacuolar membranes in contact with the void zone (Figure 6C). A contact between the membrane domains accompanied by protein separation has been detected in the nucleus-vacuole junction (NVJ), which is one of the MCSs that excludes certain proteins from the nuclear and vacuolar membranes (Pan *et al*., 2000; Dawaliby and Mayer, 2010; Toulmay and Prinz, 2013). The transmembrane proteins appear to be completely excluded from the void zone (Figure 1H); hence, the lipid-protein interactions via protein tethering may be present in the V-V contact similar to that detected in the ER-PM contact sites.

## Materials and Methods

### Chemicals, Media, and Genetic Manipulation

Chemicals were purchased from Wako Pure Chemicals Industries (Osaka, Japan) unless indicated otherwise. Standard genetic manipulations and plasmid transformation of yeast were performed as described previously (Elble, 1992; Guthrie and Fink, 2002). Yeast strains were cultured in rich YPDA medium containing 1% yeast extract (Difco Laboratories, MI, USA), 2% Bacto peptone (Difco), 2% glucose, and 0.01% adenine, or synthetic dextrose (SD) medium containing 0.17% yeast nitrogen base w/o amino acids and ammonium sulfate (Difco), 0.5% ammonium sulfate, 2% glucose, and the required amino acids or nucleic acid bases. To induce the *GAL1* promoter, 3% galactose and 0.2% sucrose were used as carbon sources instead of glucose (YPGA medium). To deplete PS in the *P_GAL1_-3HA-CHO1* background cells, the cells were grown on YPDA plates for more than 1 day before inoculation into YPDA medium. Strains carrying *URA3*-harbouring plasmids were cultured in SD medium containing 0.5% casamino acids (Difco), 0.03% tryptophan, and 0.01% adenine (SDA-U). When *cho1*Δ or Cho1-depleted strains were cultured in SD or SDA-U medium, ethanolamine was added to the final concentration of 1 mM. For serial dilution spot assays, cells were grown to early log phase in an appropriate medium and adjusted to a concentration of 1.0 × 10^7^ cells/ml. After serial tenfold dilution, 4-μl drops were spotted onto appropriate plates.

### Strains and Plasmids

Yeast strains and plasmids used in this study are listed in Tables S1 and S2, respectively. Standard molecular biological techniques were used for the construction of the plasmids, PCR amplification, and DNA sequencing (Sambrook and Russell, 2001). PCR-based procedures were used to construct the gene deletions and gene fusions with GFP, mRFP, 3HA, and the *GAL1* promoter (Longtine *et al*., 1998). All constructs produced by the PCR-based procedure were verified by colony PCR to confirm that the replacement occurred at the expected locus. Sequences of the PCR primers are available upon request.

To construct the *LEU2::mRFP-GAS1* strain, pMF608 was linearized and integrated into the *LEU2* locus. To construct the pRS316-mRFP-GAS1 (pKT2191), the SacII-XmaI fragment from pMF608 was inserted into the SacII-SmaI gap of pRS316. To construct pRS416-GFP-GAP1C (pKT2205), the C-terminal cytoplasmic region of Gap1 corresponding to 552-602 amino acids (Popov-Čeleketić *et al*., 2016) and 257 bp downstream of the *GAP1* coding region were amplified by PCR, and *PEP12* region of pRS416-GFP-PEP12 (pKT1487) (Furuta *et al*., 2007) was replaced with the PCR product. To construct pRS316-mCherry-SPO20PABD (pKT2206), the DNA fragments of mCherry and the PA-binding region of Spo20 corresponding to 51-91 amino acids (Nakanishi *et al*., 2004) were inserted into pRS316 along with *P_TPI1_* and *T_ADH1_*.

### Microscopic Observations

Cells were observed using a Nikon ECLIPSE E800 microscope (Nikon Instec, Tokyo, Japan) equipped with an HB-10103AF super-high-pressure mercury lamp and a 1.4 numerical aperture 100× Plan Apo oil immersion objective lens (Nikon Instec) with appropriate fluorescence filter sets (Nikon Instec) or differential interference contrast optics. Images were acquired using a cooled digital charge-coupled device (CCD) camera (C4742-95-12NR; Hamamatsu Photonics, Hamamatsu, Japan) and the AQUACOSMOS software (Hamamatsu Photonics).

GFP-, mRFP-, or mCherry-tagged proteins were observed in the living cells grown in early to mid-logarithmic phase, harvested, and resuspended in SD medium. Cells were immediately observed using a GFP bandpass (for GFP) or a G2-A (for mRFP and mCherry) filter set. Observations were compiled based on the examination of at least 100 cells. For supplementation of 18:1 lyso-PS (Sigma-Aldrich, St. Louis, MO, USA), a stock solution (10 mg/mL in 0.1% Nonidet P-40) was added to the culture media to the final concentration of 20 μM. For sterol staining, cells were fixed with 5% formaldehyde, washed with PBS, and labelled for 10 min in 0.5 mg/ml filipin (Sigma-Aldrich) in PBS. For staining of the plasma membrane by FM4-64 (Thermo Fisher Scientific, MA, USA), cell suspensions were mixed with an equal volume of 100 μM FM4-64 on a glass slide and observed immediately. For staining of the vacuolar membranes, cells were labelled for 15 min in 5μM FM4-64, washed with SD medium, and immediately observed. Fluorescence of filipin and FM4-64 was observed using a UV and a G2-A filter set, respectively. For the time-lapse imaging, cell suspension was spotted onto a thin layer of SD medium containing 1 mM ethanolamine and 2% agarose on a glass slide, which was quickly covered with a coverslip. During the time-lapse imaging, the sample was maintained at 37°C by a Thermo Plate (Tokai Hit, Fujinomiya, Japan). The incidence of the void zone was assessed based on three or more images of different focal planes, and based on single focal plane in the time-lapse imaging.

### Freeze-fracture replica labelling

Yeast cells sandwiched between a 20-μm–thick copper foil and a flat aluminium disc (Engineering Office M. Wohlwend, Sennwald, Switzerland) were quick-frozen by high-pressure freezing using an HPM 010 high-pressure freezing machine according to the manufacture’s instruction (Leica Microsystems, Wetzlar, Germany). The frozen specimens were transferred to the cold stage of a Balzers BAF 400 apparatus and fractured at −115° to −105°C under vacuum at ~ 1 × 10^−6^ mbar.

For genuine morphological observation, samples were exposed to electron-beam evaporation of platinum-carbon (Pt/C) (1–2 nm thickness) at an angle of 45° to the specimen surface followed by carbon (C) (10–20 nm) at an angle of 90°. After thawing, the replicas were treated with household bleach to digest biological materials before mounting on the EM grids for observation.

For labelling, freeze-fractured samples were subjected to a three-step electron-beam evaporation: C (2–5 nm), Pt/C (1–2 nm), and C (10–20 nm) as described previously (Fujita *et al*., 2010). Thawed replicas were treated with 2.5% SDS in 0.1 M Tris-HCl (pH 8.0) at 60°C overnight; with 0.1% Westase (Takara Bio) in McIlvain buffer (37 mM citrate and 126 mM disodium hydrogen phosphate, pH 6.0) containing 10 mM EDTA, 30% fetal calf serum, and a protease inhibitor cocktail for 30 min at 30°C; and with 2.5% SDS in 0.1 M Tris-HCl (pH 8.0) at 60°C overnight. For labelling of GFP-Pma1, replicas were incubated at 4°C overnight with a rabbit anti-GFP antibody in PBS containing 1% BSA followed by colloidal gold (10 nm)-conjugated protein A (University of Utrecht, Utrecht, The Netherlands) for 60 min at 37°C in 1% BSA in PBS. Replicas were observed and imaged with a JEOL JEM-1011 EM (Tokyo, Japan) using a CCD camera (Gatan, Pleasanton, CA, USA).

### Lipid analysis

Total lipids were extracted basically by the Bligh and Dyer method (Bligh and Dyer, 1959). Cells were grown in 100-200 ml of YPDA medium to 0.8-1.0 OD_600_/ml at 30°C or 37°C. The cells were collected and resuspended in 3.8 ml of chloroform-methanol-0.1 M HCl/0.1 M KCl (1:2:0.8) and lysed by vortexing with glass beads for 1 min. Then, 1.0 ml each of chloroform and 0.1 M HCl/0.1 M KCl were added followed by centrifugation, isolation of the lipid-containing phase, and evaporation of the solvent. The extracted lipids were dissolved in an appropriate volume of chloroform. Total phospholipids were determined by the phosphorus assay (Rouser *et al*., 1970).

For ergosterol analysis, the samples containing 20 nmol phosphate were subjected to TLC on an HPTLC plate (Silica gel 60; Merck Millipore, MA, USA) in the solvent system hexane/diethyl ether/acetic acid (80:20:1) (Dodge and Phillips, 1967). After migration, the plates were dried and sprayed with a 10% (w/v) cupric sulfate solution in 8% (w/v) orthophosphoric acid. Plates were heated in an oven at 180°C for 20 min. Plates were scanned with a CanoScan 8800F image scanner (Canon, Tokyo, Japan) and the acquired images were quantified using the ImageJ software.

### Statistical Analysis

Significant differences in Figures 5C and 7C were determined using a two-sided Student’s *t* test. Significant differences for all other figures were determined by the Tukey–Kramer test.

## Acknowledgements

We thank Masahiko Watanabe (Hokkaido University, Sapporo, Japan) for providing the rabbit anti-GFP antibody for freeze-fracture replica labelling. We thank Takehiko Yoko-o (AIST, Tokyo, Japan) for providing the pMF608 plasmid. We thank our colleagues in the Tanaka laboratory for valuable discussions and Eriko Itoh for technical assistance. This work was supported by Japan Society for the Promotion of Science (JSPS) KAKENHI grants 18K14645 (T.M.), 18K06104 (T.K.), and 19K06536 (K.T.). This work was partly supported by the Photo-excitonix Project at Hokkaido University.

