## Supplemental files for "Phosphatidylserine prevents the generation of a protein-free giant plasma membrane domain in yeast"

**Figure S1**

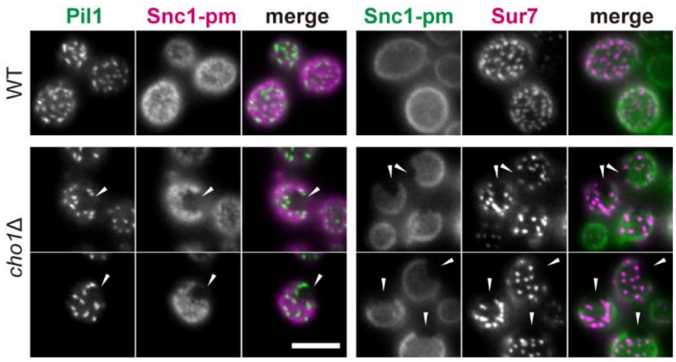


**Figure S1 Eisosome structures are excluded from the void zone**

Eisosome components were not present in the void zone. Cells expressing Pil1-GFP and mRFP-Snc1-pm or GFP-Snc1-pm and Sur7-mRFP were grown in YPDA medium at 37°C. Images shown are focused on the cell surface. Arrowheads indicate the void zone. Scale bar: 5 μm.

**Figure S2**

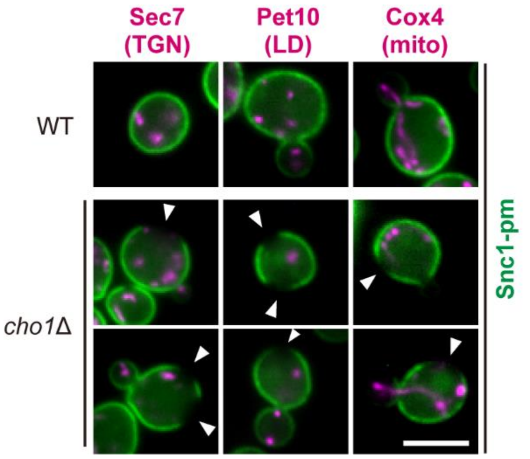


**Figure S2 Distribution of organelles in the cells with the void zone.**

The distribution of the *trans*-Golgi network (TGN), lipid droplets (LD), and mitochondria (mito) is not influenced by the void zone. Cells expressing GFP-Snc1-pm and Sec7-mRFP, Pet10-GFP and mRFP-Snc1-pm, or Cox4-GFP and mRFP-Snc1-pm were grown in YPDA or SDA-U media at 37°C. Snc1-pm and the organelle marker proteins are shown in green and magenta, respectively. Arrowheads indicate the void zone. Scale bar: 5 µm.

**Figure S3**

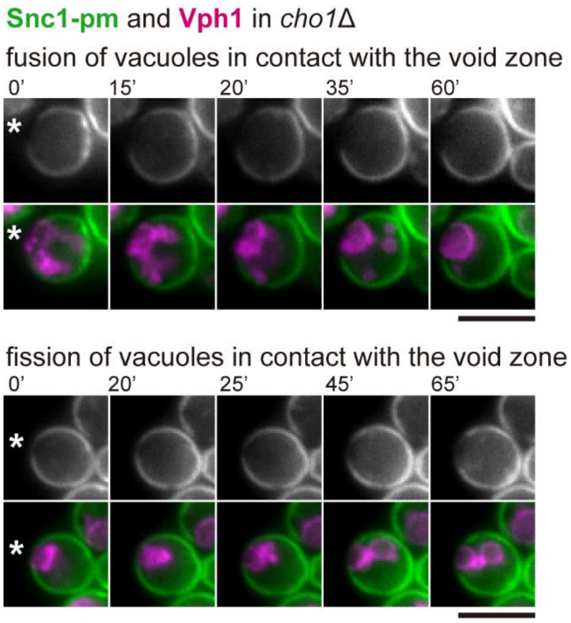


**Figure S3 Fusion and fission of vacuoles in contact with the void zone.**

Time-lapse imaging of the vacuole-void zone contact was performed. *cho1*Δ cells expressing GFP-Snc1-pm and Vph1-mRFP were grown in YPDA medium at 37°C. Fusion and fission of vacuoles in contact with the void zone are shown in the upper and lower panels, respectively. Numbers indicate time in min. Asterisks indicate the V-V contact. Scale bars: 5 µm.

**Figure S4**

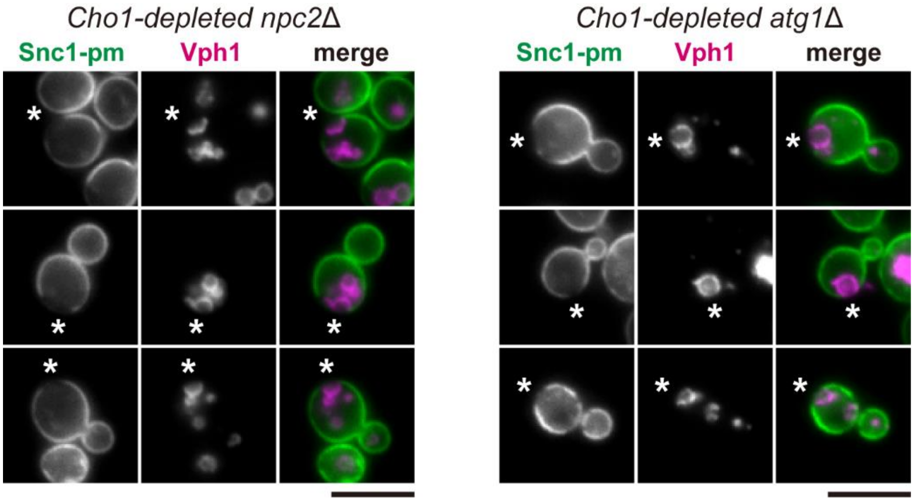


**Figure S4 Npc2 and Atg1 are not essential for the formation of the void zone and the V-V contacts.**

The void zone and the V-V contact in the absence of Npc2 and Atg1. Cells expressing GFP-Snc1-pm and Vph1-mRFP were grown in YPDA medium at 37°C. Asterisks indicate the V-V contact. Scale bars: 5 µm.

**Movie 1 (thumbnail image)**

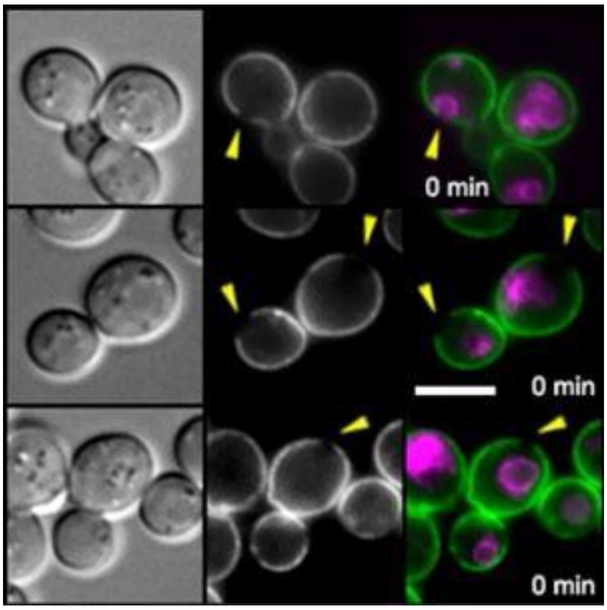


**Movie 1**

Time-lapse imaging of the vacuole-void zone contact. Time-lapse imaging was performed as in Figure 6D and Materials and Methods. Three samples are shown. Yellow arrowheads indicate the void zone. Yellow circles indicate the V-V contact site. Scale bar: 5 µm.
